# Transient upregulation of procaspase-3 during oligodendrocyte fate decisions

**DOI:** 10.1101/2024.11.13.623446

**Authors:** Yasmine Kamen, Timothy W. Chapman, Enrique T. Piedra, Matthew E. Ciolkowski, Robert A. Hill

## Abstract

Oligodendrocytes are generated throughout life and in neurodegenerative conditions from brain resident oligodendrocyte precursor cells (OPCs). The transition from OPC to oligodendrocyte involves a complex cascade of molecular and morphological states that position the cell to make a fate decision to integrate as a myelinating oligodendrocyte or die through apoptosis. Oligodendrocyte maturation impacts the cell death mechanisms that occur in degenerative conditions, but it is unclear if and how the cell death machinery changes as OPCs transition into oligodendrocytes. Here, we discovered that differentiating oligodendrocytes transiently upregulate the zymogen procaspase-3, equipping these cells to make a survival decision during differentiation. Pharmacological inhibition of caspase-3 decreases oligodendrocyte density, indicating that procaspase-3 upregulation promotes differentiation. Moreover, using procaspase-3 as a marker, we show that oligodendrocyte differentiation continues in the aging cortex and white matter. Taken together, our data establish procaspase-3 as a differentiating oligodendrocyte marker and provide insight into the underlying mechanisms occurring during the decision to integrate or die.

## INTRODUCTION

Myelination continues into late adulthood and is thought to contribute to learning and memory (Fields, 2010; Hill et al., 2018; Hughes et al., 2018; McKenzie et al., 2014; Pan et al., 2020; Shimizu et al., 2023; Steadman et al., 2020; Xiao et al., 2016). Myelin is produced by oligodendrocytes, which arise from brain resident oligodendrocyte precursor cells (OPCs) through a multi-step differentiation process characterized by distinct morphological, molecular, and metabolic changes (Emery, 2010; Bergles and Richardson, 2015; Hill et al., 2024). The cell stages occurring during differentiation can be roughly separated into premyelinating oligodendrocytes, newly formed oligodendrocytes, and mature oligodendrocytes (Marques et al., 2016; Stadelmann et al., 2019). However, the precise definition and delineation of these stages depends on the approaches used and remains a challenge for the field. This is important because these stages are thought to represent critical cellular checkpoints that result in either the successful or the failed integration of mature myelinating oligodendrocytes that will likely remain in the brain for life.

The cell checkpoints that result in successful oligodendrocyte generation are also central for understanding myelin pathology and ways to treat it. Myelin dysfunction and oligodendrocyte death occur in several neurodegenerative disorders such as Multiple Sclerosis and Alzheimer’s disease (Franklin, 2002; Calabrese et al., 2015; Chen et al., 2021; Depp et al., 2023). Moreover, myelin loss is prominent in aging and is thought to contribute to cognitive decline (Peters et al., 1996; Peters and Kemper, 2012; Wang et al., 2020). Understanding the mechanisms underlying these checkpoints could therefore uncover ways to promote the generation of oligodendrocytes to replace the lost myelin.

Oligodendrocyte maturation impacts the vulnerability and cell death mechanisms following different insults (Back et al., 1998, 2001; Khorchid et al., 2002; Baerwald and Popko, 1998; Chapman et al., 2024). For instance, treatment of cultured cells with interferon-γ induces apoptosis in developing oligodendrocytes, but causes necrosis in mature oligodendrocytes, both occurring over different time courses (Baerwald and Popko, 1998). Similarly, in vivo single cell phototoxic DNA damage or cuprizone-mediated demyelination result in different cell death time courses for premyelinating and newly formed oligodendrocytes compared to mature oligodendrocytes, reflecting different cell death mechanisms across the lineage (Chapman et al., 2023, 2024).

Death of differentiating oligodendrocytes is also a part of normal development, as previous reports suggest that 20-40% of cortical premyelinating oligodendrocytes die during the first postnatal month, a proportion that increases to 80% in middle-aged mice (Trapp et al., 1997; Hughes et al., 2018). This has been proposed to regulate the spatial and temporal specificity of developmental myelination, as removal of apoptotic machinery in the oligodendrocyte lineage during early postnatal development leads to early and ectopic myelination (Sun et al., 2018). In contrast, mature oligodendrocytes are long-lived and are rarely observed dying in young and middle-aged mice (Tripathi et al., 2017; Hill et al., 2018; Hughes et al., 2018; Chapman et al., 2023). This raises the question of whether oligodendrocyte maturation alters apoptotic machinery and impacts the death mechanism or the propensity to die, not only during disease associated degenerative conditions, but also during normal development and aging.

Here, we discovered that oligodendrocytes upregulate the zymogen procaspase-3 specifically during the premyelinating and newly formed stages, suggesting that they are primed to make a fate decision between cell death or differentiation. We further find that blocking caspase-3 reduces oligodendrocyte density suggesting that this upregulation plays a role in differentiation. Taken together, our findings establish procaspase-3 as a new marker for differentiating oligodendrocytes and uncover additional underlying mechanisms occurring during oligodendrogenesis.

## RESULTS

### Procaspase-3 is upregulated in differentiating oligodendrocytes

Previous reports suggest that differentiating, but not mature, oligodendrocytes die through cleaved caspase-3-mediated apoptosis in response to cuprizone toxicity (Chapman et al., 2024), and cleaved caspase-3 labeling is observed in differentiating oligodendrocytes during early postnatal development (Sun et al., 2018). Therefore, to investigate whether oligodendrocyte maturation coincides with changes in apoptotic machinery, we examined whether the zymogen procaspase-3 differed with maturation. To do so, we co-labeled somatosensory cortical sections in 8-week-old mice with antibodies against CNP (a newly formed and mature oligodendrocyte marker (Butt et al., 1995; Vogel et al., 1988)) and procaspase-3. We found a population of cells that expressed procaspase-3 significantly above background level, were sparsely distributed across the cortex, and appeared to partially overlap with CNP+ cells (Figure 1A,B). While 82.58 ± 1.51% of procaspase-3+ cells were also CNP+, these cells only represented 8.30 ± 1.41% of the CNP cell population (Figure 1C,D). To better define the identity of these procaspase-3+ cells, we labeled somatosensory cortical sections with different markers for oligodendrocyte lineage cells (Figure 2A,B). We found that all procaspase-3+ cells were positive for OLIG2 (Figure 2A,C), indicating that they belonged to the oligodendrocyte lineage (Lu et al., 2000; Zhou and Anderson, 2002). None of the procaspase-3+ cells overlapped with PDGFRA (Figure 2A,C,D), suggesting that they were not OPCs (Nishiyama et al., 1996). In contrast, all procaspase-3+ cells were positive for BCAS1 (Figure 2A,C), a differentiating oligodendrocyte marker (Fard et al., 2017). Staining with procaspase-3 and NG2 (a second OPC marker (Nishiyama et al., 1996)) indicated that 15.75 ± 5.27% of procaspase-3+ cells were labeled with NG2 (Figure 2A,C). However, all procaspase-3+ NG2+ cells were also BCAS1+ and we did not detect any procaspase-3+ NG2+ BCAS1-cells. These data are in line with previous reports of BCAS1 and NG2 overlap (Fard et al., 2017) and are consistent with procaspase-3 being upregulated in differentiating oligodendrocytes but not OPCs. We also detected a 79.37 ± 3.64% overlap between procaspase-3 and TCF7L2, an additional differentiating oligodendrocyte marker (Guo and Wang, 2023) (Figure 2A,C). Moreover, 93.99 ± 3.75% of BCAS1+ cells were procaspase-3+ (Figure 2D), suggesting that most differentiating oligodendrocytes have high levels of procaspase-3.

**Figure 1.**
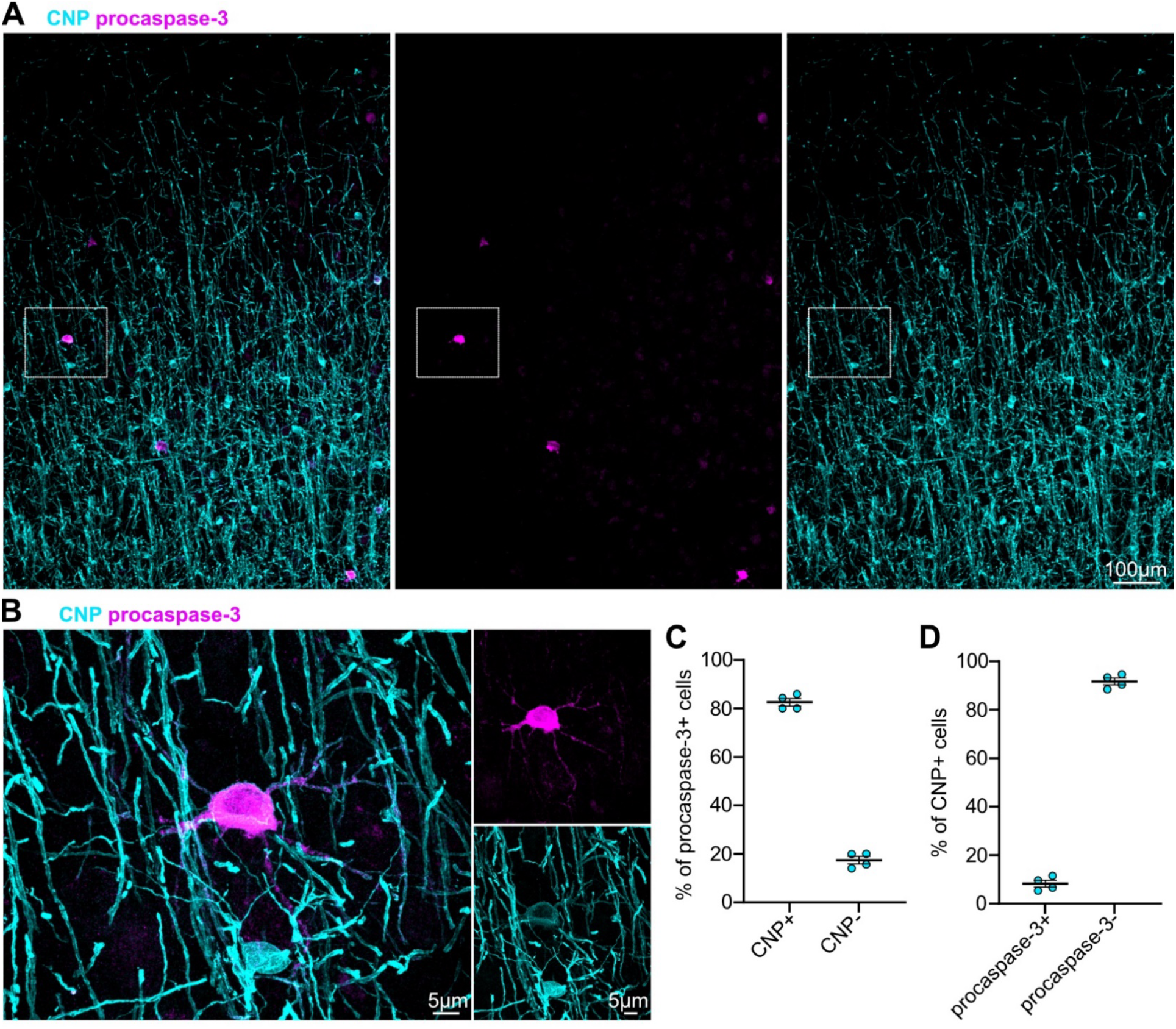
Procaspase-3 labels a subset of oligodendrocytes in the cortex. **A)** CNP and procaspase-3 staining in the somatosensory cortex of 8-week-old (P60) mice. **B)** Boxed area from **A**, with a CNP+ procaspase-3+ cell and a CNP+ procaspase-3- cell. **C)** Proportion of procaspase-3 cells that are co-labeled with CNP (*n* = 4 mice). Data are shown as mean ± s.e.m. **D)** Proportion of CNP cells that are positive for procaspase-3 (*n* = 4 mice). Data are shown as mean ± s.e.m.

**Figure 2.**
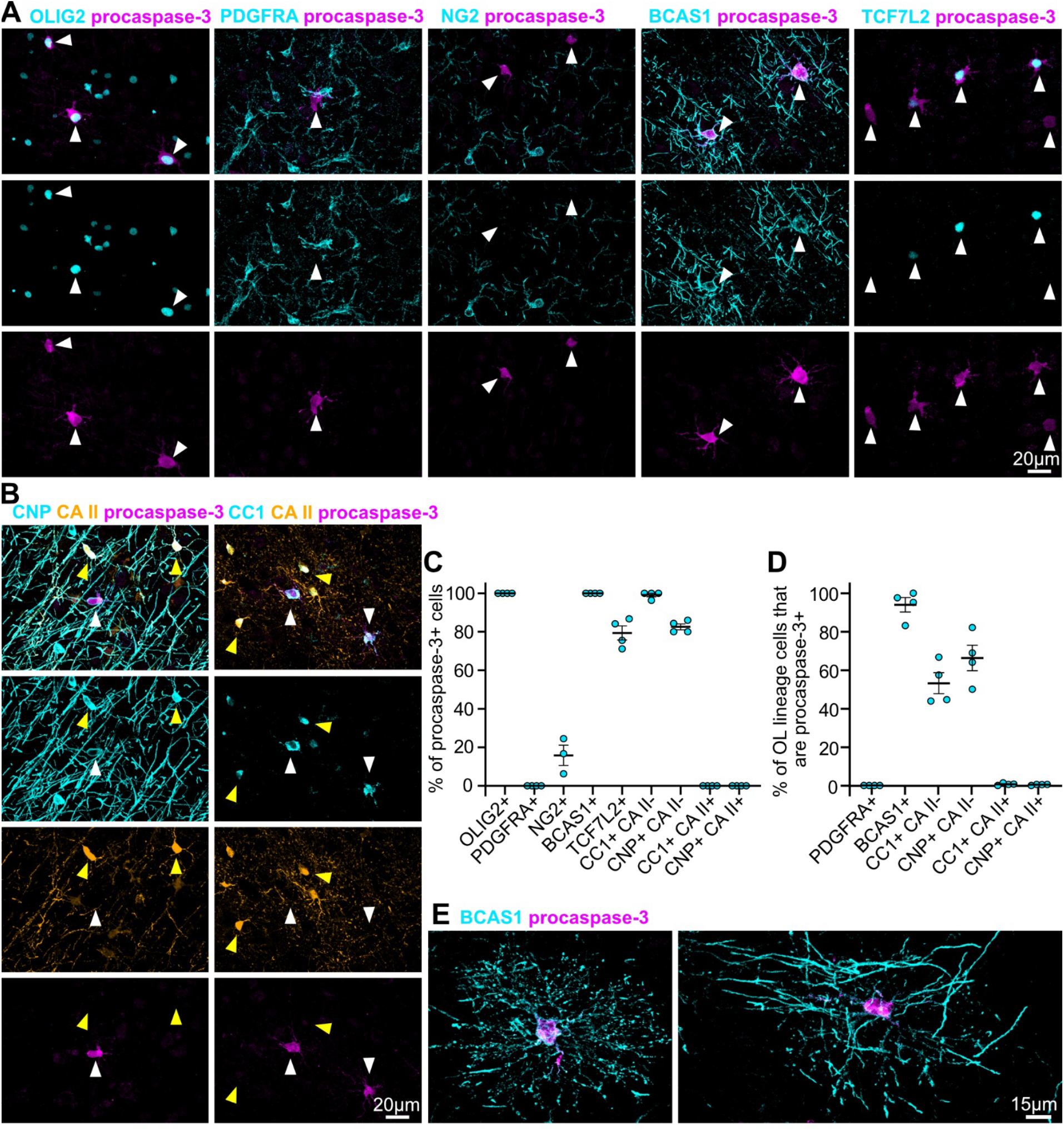
Procaspase-3 labels differentiating oligodendrocytes in the cortex. **A)** Staining against procaspase-3 and markers of different oligodendrocyte lineage stages in the cortex of 8-week-old (P60) mice. White arrowheads indicate procaspase-3+ cells. **B)** Staining against procaspase-3 and oligodendrocyte markers. White arrowheads indicate procaspase-3+ cells and yellow arrowheads indicate CA II+ procaspase-3-cells. **C)** Proportion of procaspase-3+ cells that are co-labeled by oligodendrocyte lineage markers (*n* = 4 mice). Data are shown as mean ± s.e.m. **D)** Proportion of oligodendrocyte lineage cells that are procaspase-3+ (*n* = 4 mice). Data are shown as mean ± s.e.m. **E)** Examples of procaspase-3+ oligodendrocytes without (left) or with (right) myelin sheaths.

As we detected an overlap between procaspase-3+ and CNP+ cells, but CNP labels both newly formed and mature oligodendrocytes, we next examined whether procaspase-3+ cells were confined to the differentiating oligodendrocyte stage or could also be mature oligodendrocytes. To assess this, we stained somatosensory cortical sections with procaspase-3, CNP, and carbonic anhydrase (CA II; a mature oligodendrocyte marker (Cammer, 1984; Chapman et al., 2024)), or procaspase-3, CC1 (a differentiating and mature oligodendrocyte marker (Bhat et al., 1996; Fard et al., 2017)), and CA II (Figure 2B). We found that 99.07 ± 0.93% of procaspase-3+ cells were CC1+ CA II-, and 82.58 ± 1.51% were CNP+ CA II- (Figure 1D,E). Only 0.68 ± 0.28% of CC1+CA II+ cells and 0.40 ± 0.13% of CNP+ CA II+ cells were procaspase-3+ (Figure 2D,), suggesting that procaspase-3 upregulation is confined to the differentiating stage. This is consistent with RNA sequencing data suggesting that *Casp3* is expressed at higher levels in newly formed oligodendrocytes than in OPCs or myelinating oligodendrocytes (Marques et al., 2016; Zhang et al., 2014), and procaspase-3 increasing during in vitro oligodendrocyte differentiation (Soane et al., 1999). Procaspase-3+ cells had either a premyelinating (no BCAS1+ sheaths) or newly formed oligodendrocyte (BCAS1+ sheaths) morphology (Figure 2E). Taken together, these data suggest that differentiating and mature oligodendrocytes have different pro apoptotic machinery, with differentiating oligodendrocytes transiently upregulating procaspase-3.

### Procaspase-3 labels differentiating oligodendrocytes in the corpus callosum

To examine whether procaspase-3 upregulation in differentiating oligodendrocytes was specific to the cortex or consistent across multiple brain regions, we turned to the corpus callosum of 8-week-old mice. Like in the cortex, we detected sparsely distributed cells that expressed procaspase-3 significantly above background levels (Figure 3A). To validate that these cells were differentiating oligodendrocytes, we co-labeled them with either OLIG2, PDGFRA, or CC1 and CA II (Figure 3B,C). We found that most procaspase-3+ cells (98.02 ± 1.17%) were OLIG2+ oligodendrocyte lineage cells (Figure 3B,D). The remaining procaspase-3+ OLIG2-cells might be neural progenitors adjacent to the white matter, as we detected cells with high procaspase-3 levels in germinal zones such as the subventricular zone and the dentate gyrus of the hippocampus that were not BCAS1 labeled (Figure 4).

**Figure 3.**
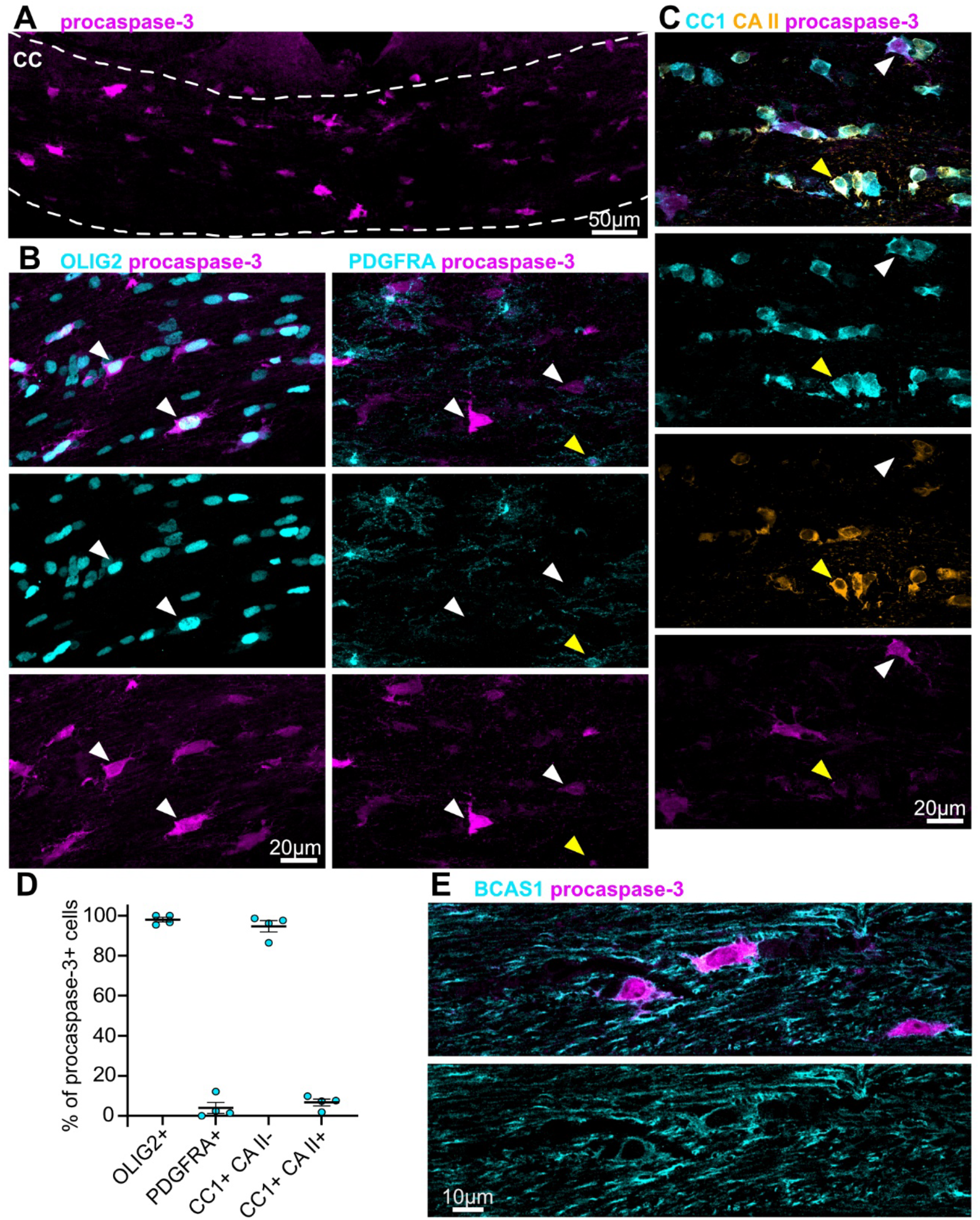
Procaspase-3 labels differentiating oligodendrocytes in the corpus callosum. **A)** Procaspase-3+ cells in the corpus callosum of 8-week-old (P60) mice. **B)** Procaspase-3 labeling along with labeling for OLIG2 and PDGFRA in the corpus callosum. The white arrow heads indicate examples of pro CASPASE3+ OLIG2+ cells or procaspase-3+ PDGFRA-cells while the yellow arrowhead indicates a procaspase-3+ PDGFRA+ cell. **C)** Procaspase-3 labeling along with labeling for CC1 and CA II. The white arrowhead indicates an example of a procaspase-3+ CC1+ CA II-cell while the yellow arrowhead indicates a procaspase-3+ CC1+ CA II+ cell. **D)** Proportion of procaspase-3+ cells that are co-labeled by oligodendrocyte lineage markers (*n* = 4 mice). Data are shown as mean ± s.e.m. **E)** Single z-plane of BCAS1 and procaspase-3 labeling in the corpus callosum.

**Figure 4.**
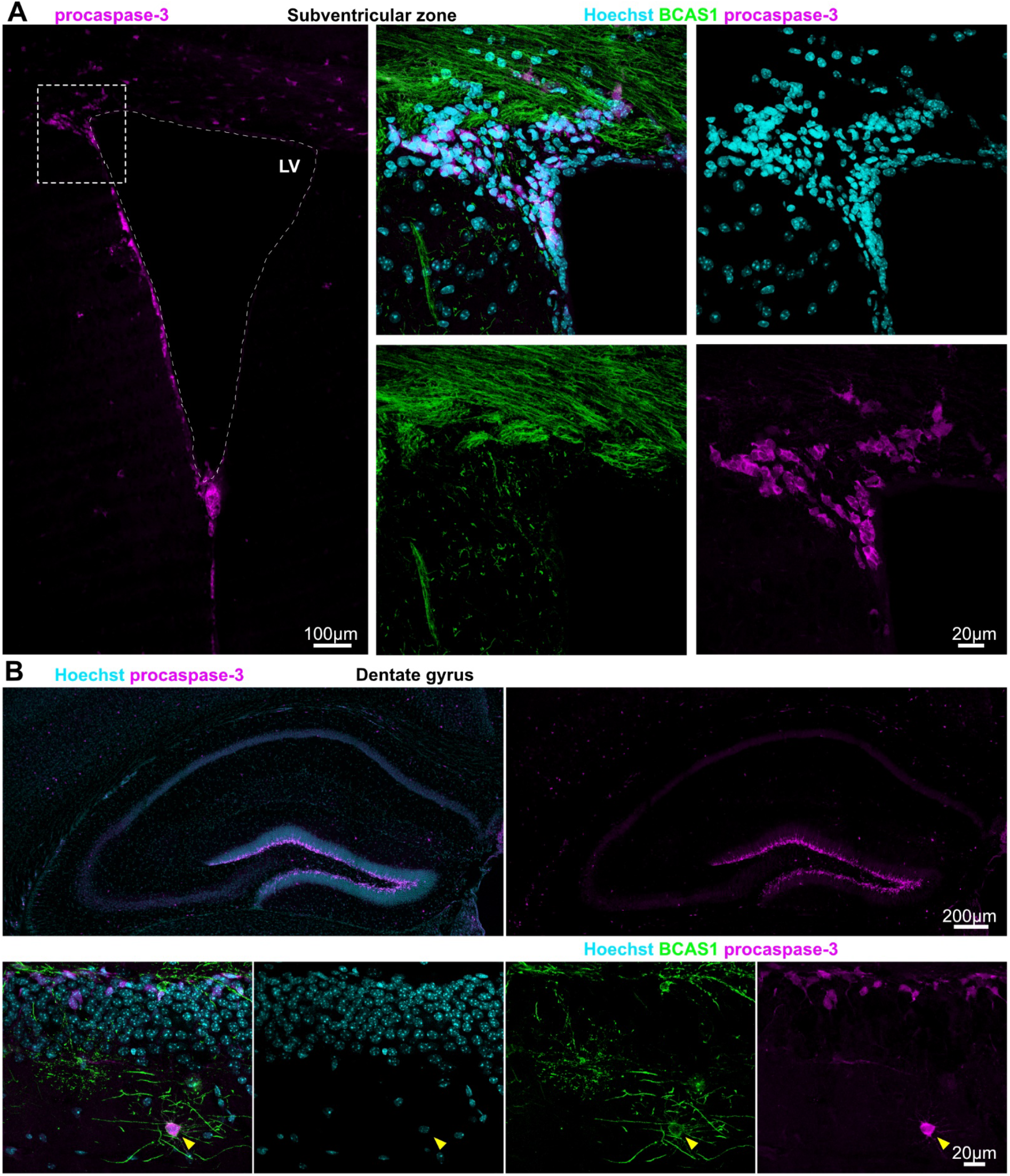
Procaspase-3 labels cells in germinal zones. **A)** BCAS1 and procaspase-3 immunolabeling in the subventricular zone of 8-week-old (P60) mice. Procaspase-3 labels a population of BCAS1-cells along the lateral ventricle wall. **B)** BCAS1 and procaspase-3 immunolabeling in the dentate gyrus of the hippocampus of 8-week-old (P60) mice. Procaspase-3 labels a population of BCAS1- cells in the dentate gyrus. The yellow arrowhead indicates a procaspase-3+ differentiating oligodendrocyte.

We only detected overlap with PDGFRA in a small proportion of procaspase-3+ cells (6.66 ± 1.7%), indicating that the majority of callosal procaspase-3+ cells were not OPCs (Figure 3B,D). As in the cortex, most procaspase-3+ cells were CC1+ CA II- (94.70 ± 2.82%), and we only detected a 3.97 ± 2.77% overlap with CC1+ CA II+ mature oligodendrocytes (Figure 3C,D). Thus, procaspase-3 is upregulated in differentiating oligodendrocytes across multiple brain regions.

BCAS1 is often used as a differentiating oligodendrocyte marker, but we found the detection of individual BCAS1+ cells in the young adult white matter difficult due to the abundance of BCAS1+ myelin (Figure 3E). As procaspase-3 appears to be cytoplasmic, it enhanced the detection of differentiating oligodendrocytes in the white matter (Figure 3E) and may therefore prove to be a useful marker for differentiating oligodendrocytes in both gray and white matter.

### Procaspase-3+ oligodendrocytes are not dying cells

Intravital imaging over several weeks in the middle-aged mouse cortex has shown that most differentiating oligodendrocytes do not stably integrate, but instead die (Hughes et al., 2018). Although 93.99 ± 3.75% of BCAS1+ differentiating oligodendrocytes in the young adult cortex have high levels of procaspase-3 and it is therefore unlikely that procaspase-3 labels dying cells, or cells destined to die, we wanted to validate that our anti-procaspase-3 antibody did not label cleaved caspase-3, to ensure that we were not labeling dying cells. To do so, we stained somatosensory cortical slices from 8-week-old mice against BCAS1, cleaved caspase-3, and procaspase-3 (Figure 5A). We found that the majority of BCAS1 cells were positive for procaspase-3 only, and not cleaved caspase-3 (92.95 ± 2.77%; Figure 5B). We only detected a small proportion of cells labeled with both procaspase-3 and cleaved caspase-3 (2.54 ± 1.41%), or with cleaved caspase-3 only (0.80 ± 0.80%) (Figure 5B). Thus, our procaspase-3 antibody does not bind cleaved caspase-3, and procaspase-3 upregulation is unlikely to indicate differentiating oligodendrocytes destined to die.

**Figure 5.**
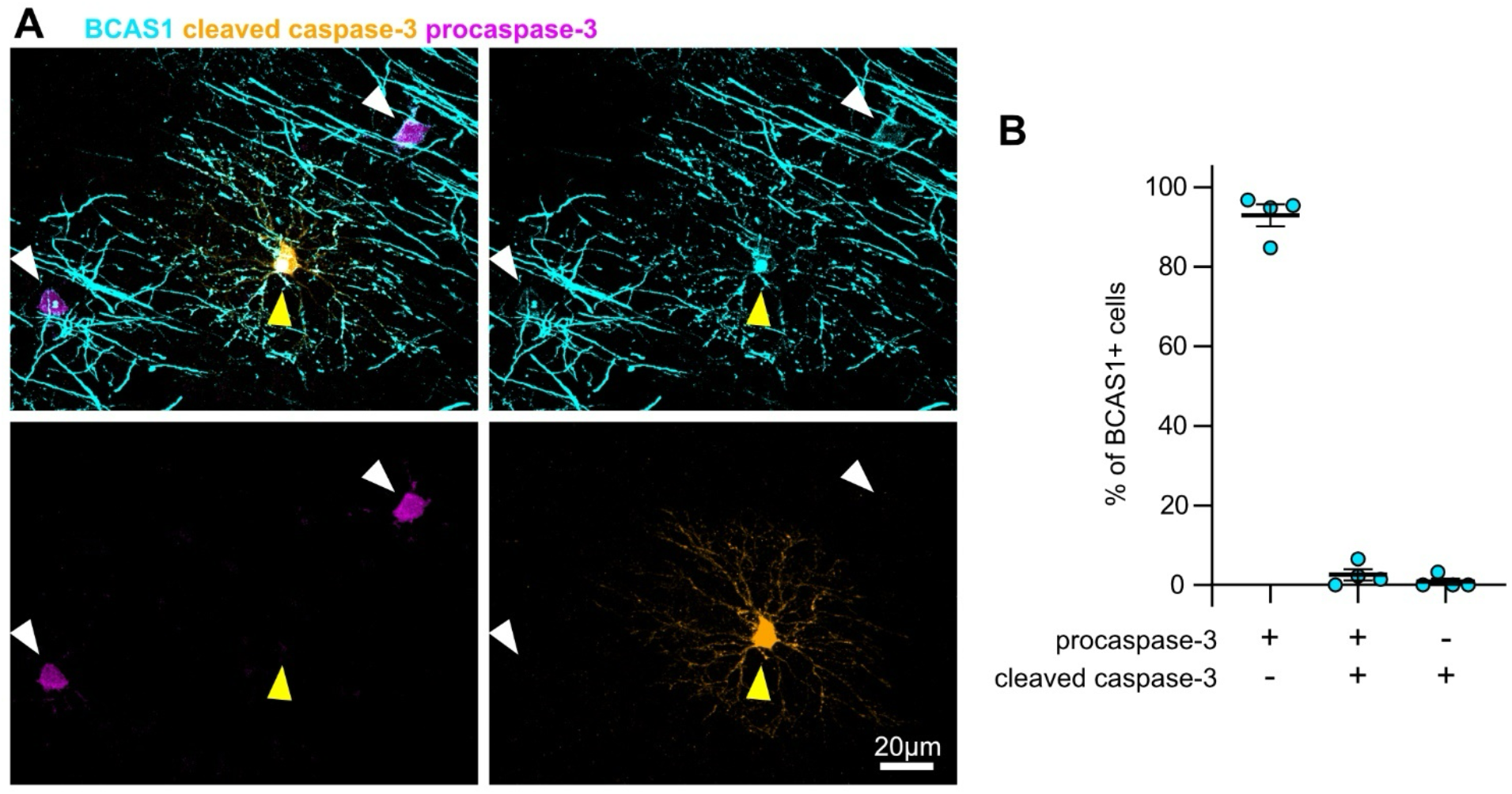
Procaspase-3 does not label dying oligodendrocytes. **A)** BCAS1, cleaved caspase-3 and procaspase-3 staining in the somatosensory cortex of 8-week-old (P60) mice indicating no overlap between cleaved caspase-3 and procaspase-3 antibodies. The white arrows indicate two BCAS1+ procaspase-3+ cleaved caspase-3-cells. The yellow arrow indicates a BCAS1+ procaspase-3-cleaved caspase-3+ cell. **B)** Proportion of BCAS1+ cells that are co-labeled with either procaspase-3 or cleaved caspase-3, or both (*n* = 4 mice). Data are shown as mean ± s.e.m.

### Procaspase3+ cell density differs with age and brain region

Oligodendrogenesis continues into adulthood, but decreases with age (Chapman et al., 2023; Hill et al., 2018; Hughes et al., 2018; Rivers et al., 2008; Young et al., 2013), and the number of BCAS1+, *Enpp6*+, or *Pcdh17it*+ (also called *lncOL1)* differentiating oligodendrocytes has been shown to decrease in middle- and old-aged mice (Fard et al., 2017; Kasuga et al., 2019). We therefore wanted to test if procaspase-3 cell density followed a similar trend. To do so, we immunolabeled the somatosensory cortex and corpus callosum of mice against procaspase-3 every three to four days during the first two postnatal weeks, monthly during the first three postnatal months, and every two to six months thereafter (Figure 6A,B). In line with our data in 8-week-old mice, across all ages 99.4% of cortical procaspase-3+ cells and 96.7% of callosal procaspase-3+ cells were positive for OLIG2 (cortex: *n* = 371 cells from 31 animals; corpus callosum: *n* = 1552 cells from 47 animals), indicating that the vast majority of procaspase-3+ cells in the cortex and corpus callosum are oligodendrocyte lineage cells throughout life.

**Figure 6.**
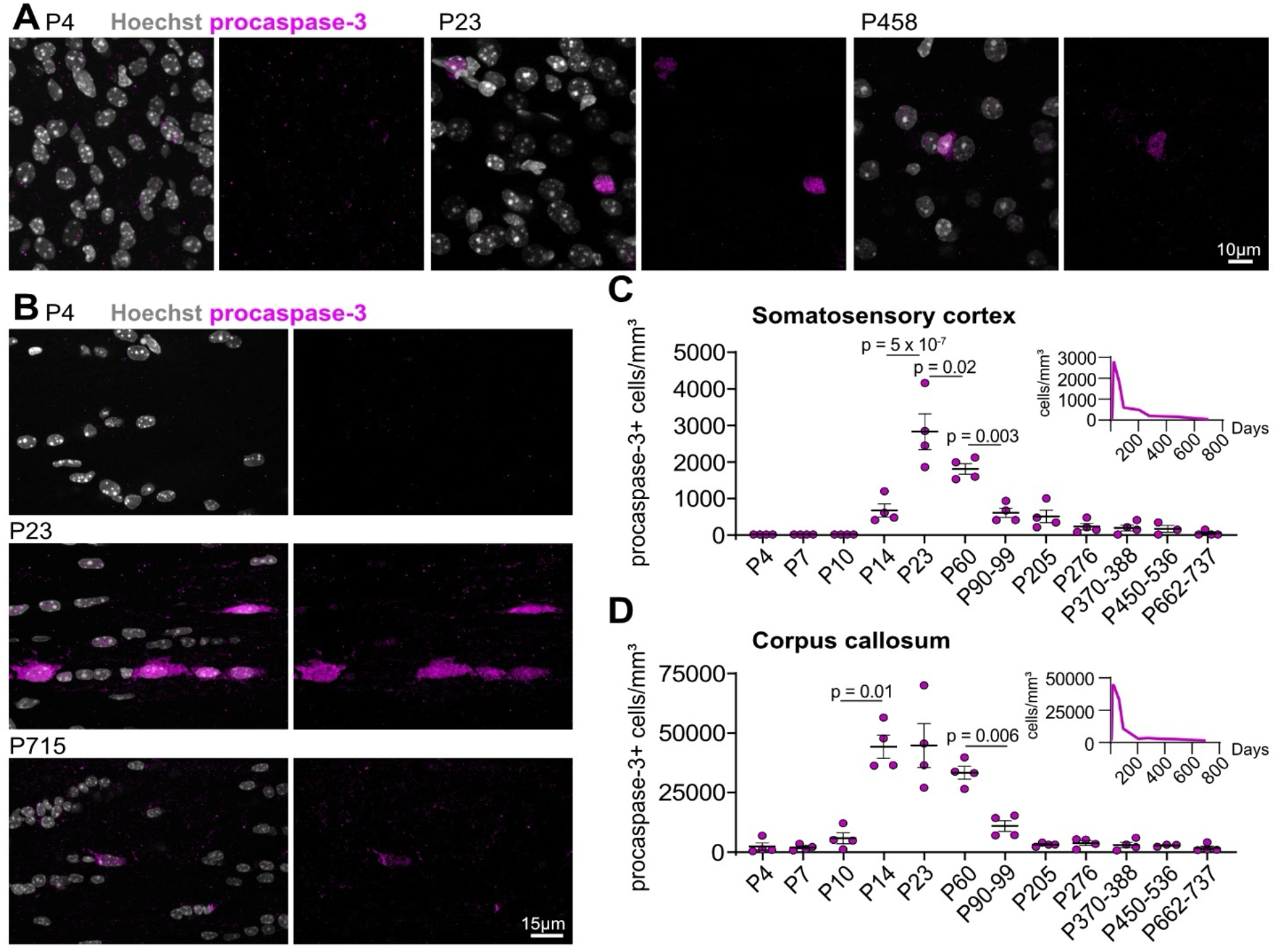
Differentiating oligodendrocyte densities differ with age and brain regions. **A)** Procaspase-3 labeling in the somatosensory cortex at P4, P23 and P458. **B)** Procaspase-3 labeling in the corpus callosum at P4, P23 and P715. **C)** Density of procaspase-3+ cells in the somatosensory cortex between P4 and P737 (*n* = 3-4 mice per timepoint). Data are shown as mean ± s.e.m. Statistical significance between P14 and P737 was assessed by one-way ANOVA (p = 2.2 × 10^−9^) and Šidák’s pairwise multiple comparisons to test for a difference between consecutive timepoints. Only p ≤ 0.05 values are displayed on the graph, and all other pairwise comparisons were p > 0.05. The inset graph displays procaspase-3+ cell density with a proportional x-axis. **D)** Density of procaspase-3+ cells in the corpus callosum between P4 and P737 (*n* = 3-4 mice per timepoint). Data are shown as mean ± s.e.m. Statistical significance between P4 and P737 was assessed with Welch’s ANOVA (p = 1.1 × 10^−6^) and Dunnett’s T3 pairwise multiple comparisons to test for a difference between consecutive timepoints. Only p ≤ 0.05 values are displayed on the graph, and all other pairwise comparisons were p > 0.05. The inset graph displays procaspase-3+ cell density with a proportional x-axis.

In the cortex, we did not detect any procaspase-3+ cells before the end of the second postnatal week (P14). Procaspase-3+ cell density then increased four-fold between P14 and P23, before decreasing at P60 and again at P90. Procaspase-3+ cell density remained stable thereafter (Figure 6A,C). These data are in line with the density of BCAS1+ oligodendrocytes decreasing after P40 (Fard et al., 2017).

In the corpus callosum, the appearance of procaspase-3+ cells occurred slightly earlier than in the cortex, as we detected procaspase-3+ cells as early as P4, although these were rare (Figure 6B,D). In addition, the number of procaspase-3+ cells appeared to reach its peak by P14 and this peak density was maintained until P60 (Figure 4D). We observed a similar trend in the adult corpus callosum as in the adult cortex, with a three-fold decrease in procaspase-3+ cell density between P60 and P90, after which densities remained similar. Of note, callosal procaspase-3+ cell densities were higher than cortical procaspase-3+ cell densities at all ages (*e*.*g*. 67-fold higher at P14, 16-fold higher at P23, and 47-fold higher at P715), in line with a higher rate of differentiation in the white matter (Rivers et al., 2008; Thornton et al., 2024; Young et al., 2013). Thus, these data suggest that oligodendrogenesis peaks in juvenile mice and continues at a reduced rate during adulthood and into aging.

### Differentiating oligodendrocytes lack procaspase-3 during early postnatal development

We were surprised to detect no cortical and very few callosal procaspase-3+ cells at P4, P7 and P10, as the presence of CC1+ cells in the cortex and corpus callosum of P6 mice, or MBP+ cells in the P7 corpus callosum has been reported before (Vincze et al., 2008; Zhu et al., 2011), and myelin can be detected in the P10 corpus callosum (Vincze et al., 2008). To determine whether the absence of procaspase-3+ cells in the cortex and corpus callosum indicated a lack of detectable differentiating oligodendrocytes in neonatal mice compared to juvenile, adult, and aged mice, we stained sections against both BCAS1 and procaspase-3 between P4 and P737 in the cortex, and P4 to P10 in the corpus callosum (Figure 7A-D). In juvenile, adult, and aged mice, the density of BCAS1+ cells followed a similar pattern as procaspase-3+ cell density in the cortex, with the majority of BCAS1+ cells also procaspase-3+ (Figure 7B). In contrast, while we detected BCAS1+ cells in the neonatal cortex and corpus callosum, we found that these lacked procaspase-3, suggesting that neonatal BCAS1+ cells do not upregulate procaspase-3 (Figure 7A-D). In the corpus callosum, a small proportion of BCAS1+ cells did have procaspase-3, but they remained a minority; in the cortex, we first detected BCAS1+ procaspase-3+ cells at P14 (Figure 7A), and this proportion increased to be the majority of BCAS1 cells by P23 (Figure 7B).

**Figure 7.**
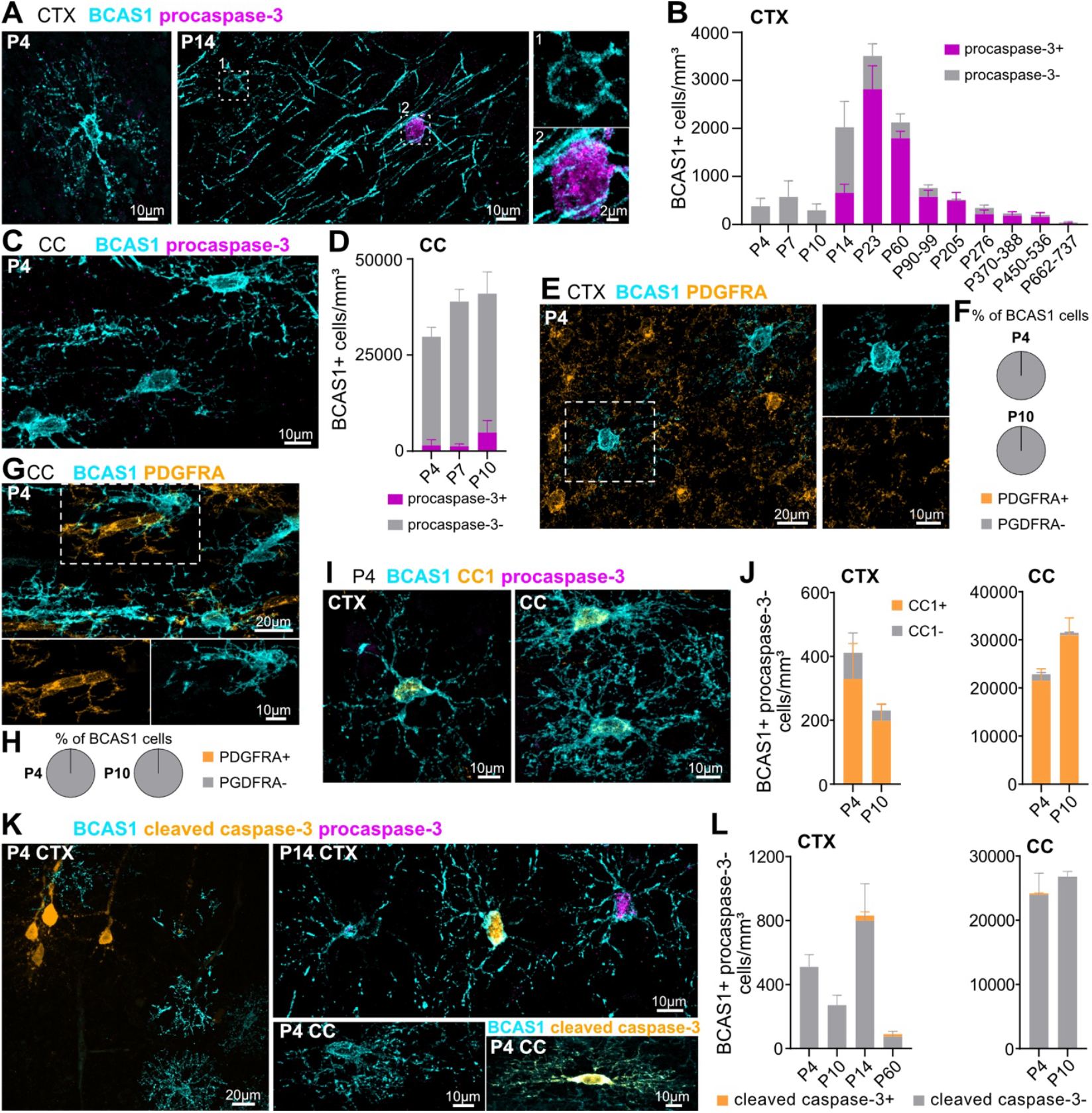
Differentiating oligodendrocytes lack procaspase-3 during early postnatal development. **A)** BCAS1 and procaspase-3 staining in the somatosensory cortex (CTX) at P4 and P14. Differentiating oligodendrocytes (BCAS1+) lack procaspase-3 at P4, and procaspase-3 labels some, but not all, differentiating oligodendrocytes at P14. **B)** BCAS1+ procaspase-3+ and BCAS1+ procaspase-3-cell density in the somatosensory cortex between P4 and P737 (*n* = 3-4 mice per timepoint). Data are shown as mean ± s.e.m. **C)** BCAS1 and procaspase-3 staining in the corpus callosum (CC) at P4. Differentiating oligodendrocytes (BCAS1+) lack procaspase-3. **D)** Density of BCAS1+ procaspase-3+ and BCAS1+ procaspase-3-cells in the corpus callosum between P4 and P10 (*n* = 3-4 mice per timepoint). Data are shown as mean ± s.e.m. **E)** BCAS1 and PDGFRA staining in the somatosensory cortex at P4, showing no overlap between cells labeled by these antibodies. **F)** The proportion of BCAS1+ cells co-labeled with PDGFRA in the somatosensory cortex at P4 and P10 (*n* = 56 cells from 4 mice for each timepoint). **G)** BCAS1 and PDGFRA staining in the corpus callosum at P4, showing no overlap between cells labeled by these antibodies. **H)** The proportion of BCAS1+ cells co-labeled with PDGFRA in the corpus callosum at P4 and P10 (*n* = 120 cells from 3 mice at P4 and *n* = 160 cells from 4 mice at P10). **I)** BCAS1 and CC1 staining in the somatosensory cortex and corpus callosum at P4, showing overlap between both markers. **J)** Density of BCAS1+ CC1+ and BCAS1+ CC1-cells in the somatosensory cortex and corpus callosum at P4 and P10 (*n* = 3-4 mice per timepoint). Data are shown as mean ± s.e.m. **K)** BCAS1, cleaved caspase-3, and procaspase-3 staining in the somatosensory cortex at P4 and P14, and in the corpus callosum at P4. The cell in the bottom right image is labeled with BCAS1 and cleaved caspase-3 only. **L)** Density of BCAS1+ procaspase-3-cleaved caspase-3+ and BCAS1+ procaspase-3-cleaved caspase-3-cells in the somatosensory cortex between P4 and P60, and in the corpus callosum at P4 and P10 (*n* = 3-4 mice per timepoint). Data are shown as mean ± s.e.m.

We next wondered if these neonatal BCAS1+ cells lacked procaspase-3 because they were still OPCs. To test this, we stained slices from P4 and P10 mice with BCAS1 and PDGFRA and analyzed whether we could detect BCAS1+ cells co-labeled with PDGFRA (Figure 7E,G). We did not detect any BCAS1+ PDGFRA+ cells in the cortex or corpus callosum at either age (Figure 7F,H), in line with previous data from the cortex of 8-week-old mice (Chapman et al., 2024).

Our findings suggest that these neonatal BCAS1+ procaspase-3-cells are not OPCs. Nonetheless, we wondered if they were not yet differentiating oligodendrocytes. We therefore labeled them with a differentiating and mature oligodendrocyte marker, CC1 (Figure 7I). The majority of P4 and P10 BCAS1+ procaspase-3-cells were positive for CC1 in both the cortex and corpus callosum, suggesting that they are indeed *bona fide* differentiating oligodendrocytes (Figure 7J).

During central nervous system development, neurons are overproduced and undergo programmed cell death (Yamaguchi and Miura, 2015); this has also been suggested for oligodendrocytes (Barres et al., 1992; Raff et al., 1993; Trapp et al., 1997) and is thought to regulate the timing of myelination (Sun et al., 2018). As we detected little overlap between procaspase-3 and cleaved caspase-3 in the 8-week-old somatosensory cortex, we wondered if these neonatal BCAS1+ CC1+ differentiating oligodendrocytes lacked procaspase-3 because they were cleaved caspase-3+ dying cells. We therefore immunolabeled the somatosensory cortex of P4, P10, P14 and P60 mice and the corpus callosum of P4 and P10 mice against BCAS1, cleaved caspase-3, and procaspase-3 (Figure 7K). Surprisingly, we detected no BCAS1+ procaspase-3-cleaved caspase-3+ cells in the P4 and P10 cortex, and only a small number at P14 and P60, coinciding with the peak in BCAS1+ cell density (Figure 7B,K,L). Similarly, we only detected a small number of callosal BCAS1+ procaspase-3-cleaved caspase-3+ cells at P4, and none at P10 (Figure 7K,L). Thus, BCAS1+ differentiating oligodendrocytes do not appear to undergo a large caspase-3-dependent cell death event. Collectively, these data suggest that differentiating oligodendrocytes do not upregulate procaspase-3 during early postnatal development, in contrast to adulthood.

### Inhibiting caspase-3 reduces oligodendrocyte density

As procaspase-3 is upregulated specifically in differentiating oligodendrocytes, we next asked whether blocking procaspase-3 would alter differentiation. To do so, we prepared hippocampal slice cultures from P9 mice and after seven recovery days in vitro (DIV), treated the slices with a specific caspase-3 inhibitor, Z-DEVD-FMK (Kanthasamy et al., 2006), or the vehicle for a further seven days (Figure 8A,B). Importantly, this inhibitor lowers both procaspase-3 and cleaved caspase-3 protein levels (Liu et al., 2015). We found that treatment with Z-DEVD-FMK significantly reduced the density of procaspase-3+ cells and trended toward decreasing the density of cleaved caspase-3+ cells, although this did not reach statistical significance (Figure 8C-E). This reduction in procaspase-3+ cells resulted in a decrease in CNP+ cells (Figure 8F,G), suggesting that blocking caspase-3 reduces oligodendrocyte differentiation. Thus, procaspase-3 upregulation may promote oligodendrocyte differentiation.

**Figure 8.**
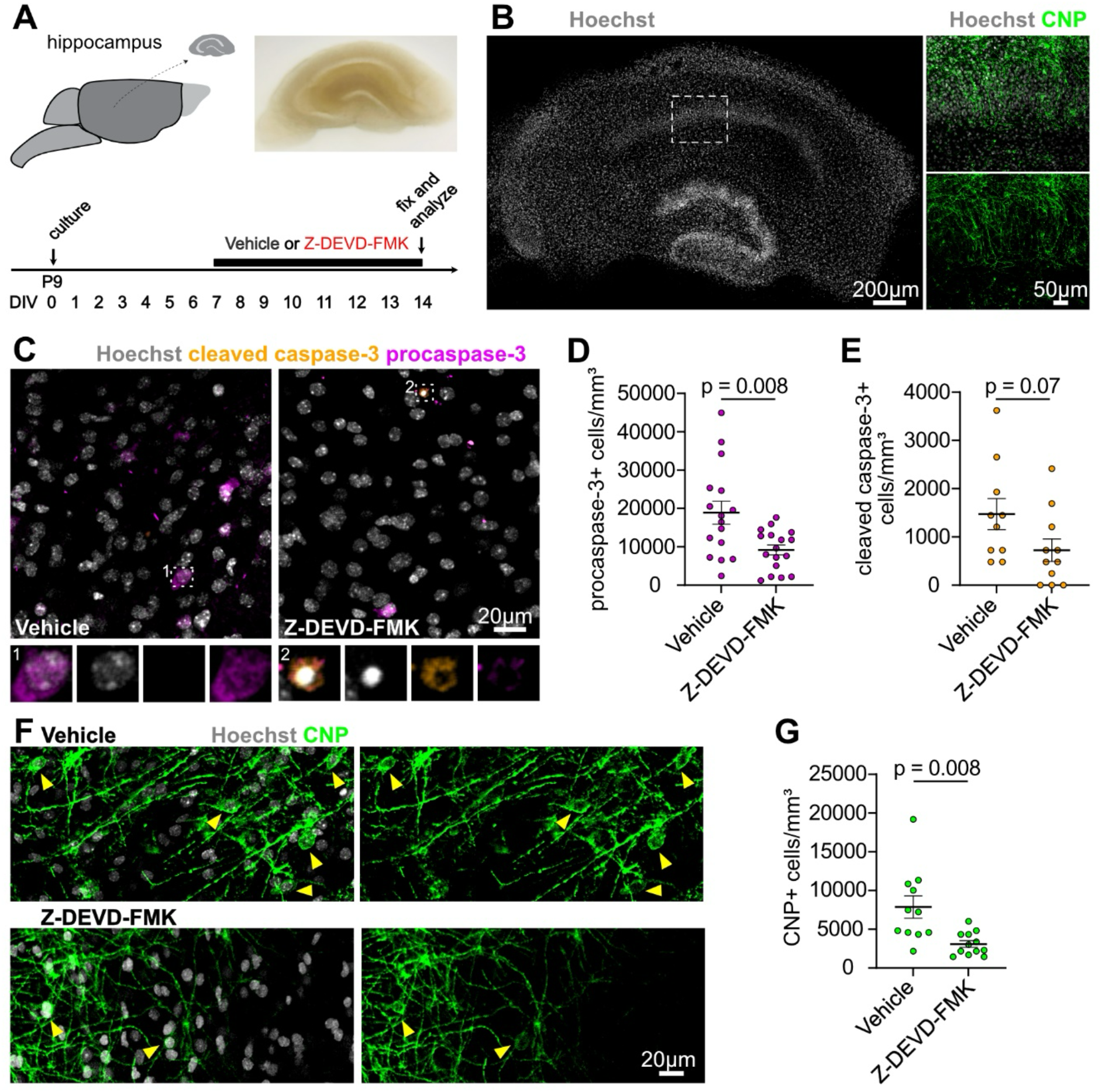
Blocking caspase-3 decreases oligodendrocyte density. **A)** Experimental design. Hippocampal slices were prepared from P9 mice and cultured for 7 days before treatment with a caspase-3 inhibitor, Z-DEVD-FMK (20 μM), or the vehicle only for a further 7 days. **B)** Hippocampal cultured slice labeled with nuclear dye. The boxed rectangle is shown on the right with nuclear dye and CNP labeling. **C)** Procaspase-3 and cleaved caspase-3 staining in the cultured slices. **D)** Density of procaspase-3+ cells in vehicle or Z-DEVD-FMK treated slices. Treatment with Z-DEVD-FMK reduced the density of procaspase-3+ cells (*n* = 16 slices from 9 mice (vehicle) or 17 slices from 9 mice (Z-DEVD-FMK) from two separate experiments; unpaired two-tailed t-test with Welch’s correction for unequal variance). Data are shown as mean ± s.e.m. **E)** Density of cleaved caspase-3+ cells in vehicle or Z-DEVD-FMK treated slices. Treatment with Z-DEVD-FMK did not significantly alter the density of cleaved caspase-3+ cells (*n* = 10 slices from 9 mice (vehicle) or 11 slices from 9 mice (Z-DEVD-FMK) from two separate experiments; unpaired two-tailed t-test). Data are shown as mean ± s.e.m. **F)** CNP staining in the cultured slices. The yellow arrow heads indicate CNP+ cells. **G)** Density of CNP+ cells in vehicle or Z-DEVD-FMK treated slices. Treatment with Z-DEVD-FMK reduced the density of CNP+ cells (*n* = 11 slices from 10 mice (vehicle) or 12 slices from 10 mice (Z-DEVD-FMK) from two separate experiments; unpaired two-tailed t-test with Welch’s correction for unequal variance). Data are shown as mean ± s.e.m.

## DISCUSSION

We set out to determine whether differentiating and mature oligodendrocytes differ in their pro apoptotic machinery. We serendipitously discovered that differentiating oligodendrocytes upregulate procaspase-3, suggesting that they are transiently primed for a fate decision between terminal differentiation and death. This also means that procaspase-3 can be used as a marker for this maturation stage (summarized in Figure 9). Proteomic and transcriptomic screens have recently allowed the identification of several new markers for differentiating oligodendrocytes, including BCAS1, ENPP6, ITPR2, or TCF7L2 (Fard et al., 2017; Guo and Wang, 2023; Marques et al., 2016; Morita et al., 2016; Xiao et al., 2016; Zhang et al., 2014), as well as *Pcdh17it/lncOL1* transcripts (He et al., 2017; Kasuga et al., 2019). Both BCAS1 and ENPP6 label myelin sheaths, making them ideal to visualize morphology of sparsely distributed cells, but limiting single-cell detection in heavily myelinated regions. In contrast, the nuclear marker TCF7L2 allows for easy single-cell detection, but does not reveal morphology, nor does in situ hybridization for *Pcdh17it/lncOL1*. Our data suggest that procaspase-3 can complement existing differentiating oligodendrocyte markers, allowing for cytoplasmic labeling that can partially reveal morphology, as well as the identification of single cells in both sparsely populated and heavily myelinated areas.

**Figure 9.**
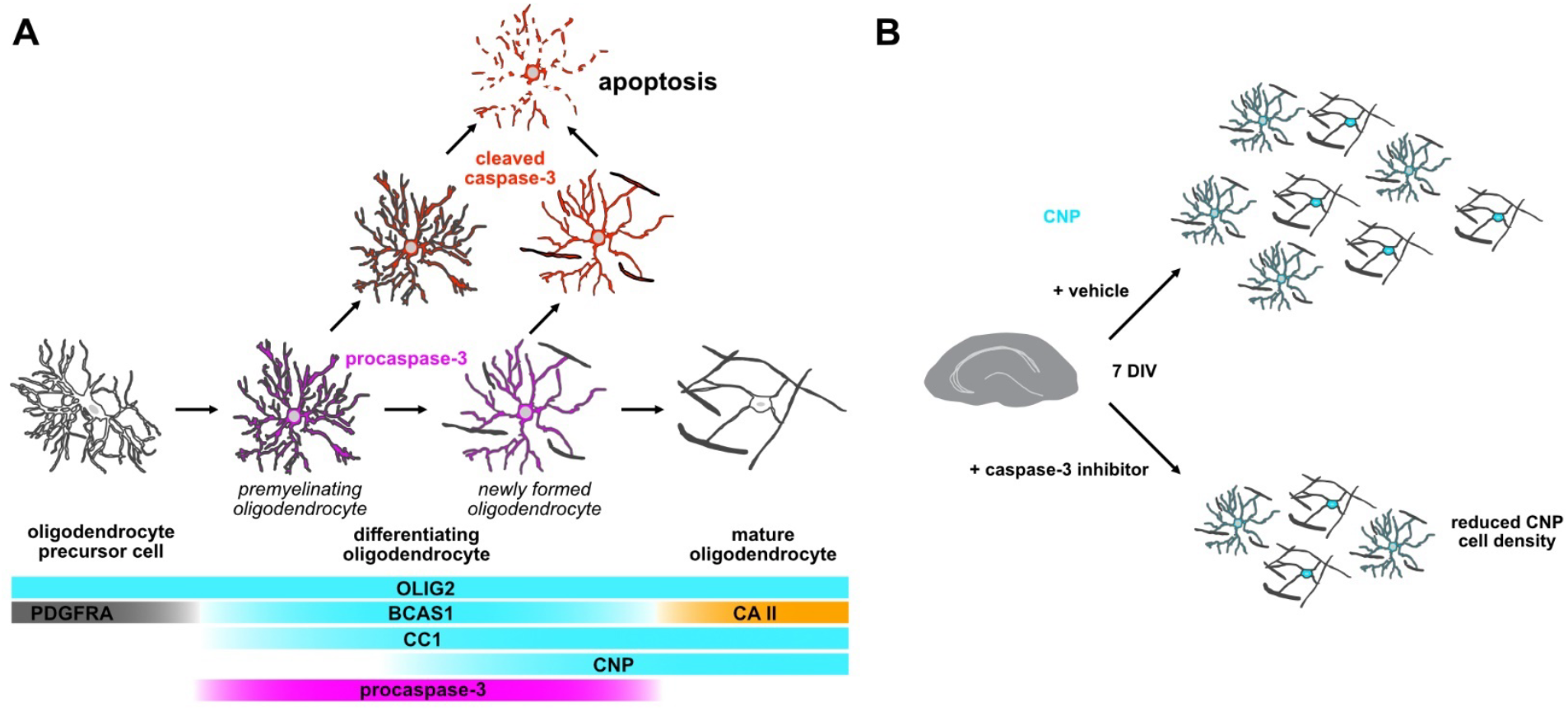
Summary of findings. **A)** Schematic with common molecular markers used to define stages in the oligodendrocyte lineage. Our data indicate that procaspase-3 labels differentiating oligodendrocytes (both premyelinating and newly formed), suggesting that these cells are primed for a fate decision between terminal differentiation into mature oligodendrocytes or cleaved caspase-3-mediated apoptosis. **B)** Treatment with a caspase-3 inhibitor reduced CNP+ cell density in hippocampal slice cultures, suggesting a role for caspase-3 in differentiation.

Using procaspase-3 as a marker of differentiating oligodendrocytes, we found that differentiating oligodendrocyte density decreased between the first, second, and third postnatal months in the cortex, and the second and third postnatal months in the corpus callosum. Nevertheless, we detected differentiating oligodendrocytes in all adult and aged cortical timepoints, consistent with previous work (Fard et al., 2017) and with protracted myelination in the cortex (Hill et al., 2018; Hughes et al., 2018). We also detected differentiating oligodendrocytes in the corpus callosum throughout adulthood, consistent with previous work demonstrating that new oligodendrocytes are generated in the 6-month-old corpus callosum (Rivers et al., 2008) or reporting a small number of *Enpp6*+ and *Pcdh17it*+ cells in the 11-month-old corpus callosum (Kasuga et al., 2019). We found that differentiating oligodendrocytes were present in the corpus callosum until at least 2 years of age, suggesting that differentiation also continues in the aged white matter. It will be interesting to test whether this is consistent in more heavily myelinated white matter tracts such as the optic nerve or spinal cord, where oligodendrogenesis has been suggested to occur until at least 1 year of age (Young et al., 2013), or in the fully myelinated brainstem.

Beyond the identification of an additional marker for differentiating oligodendrocytes, the upregulation of procaspase-3 during differentiation raises several intriguing questions, first and foremost, why is procaspase-3 upregulated? Terminal differentiation from the differentiating oligodendrocyte stage to the mature oligodendrocyte stage has been suggested to be inefficient, with 20-40% of differentiating oligodendrocytes in the developing cortex (Trapp et al., 1997), 50% of differentiating oligodendrocytes in the developing optic nerve (Barres et al., 1992), and up to 80% of differentiating oligodendrocytes in the middle-aged cortex failing to integrate and dying (Hughes et al., 2018). One possible conclusion is therefore that procaspase-3 upregulation reflects a fate decision point between death and integration. Importantly, we found that procaspase-3 did not label dying cells, indicating that procaspase-3 upregulation does not determine cell death, but rather, might be upregulated to allow for a rapid fate decision, with differentiating oligodendrocytes poised to undergo cleaved caspase-3-mediated apoptosis should they fail to receive the appropriate cues to myelinate and integrate. Intriguingly, spatial and temporal regulation of myelination has been shown to occur through the activation of the TFEB-PUMA-Bax-Bak axis, which likely induces cleaved caspase-3-mediated apoptosis of differentiating oligodendrocytes (Sun et al., 2018). Similarly to *Casp3*, both *Tfeb* and *Bbc3* transcripts are upregulated in differentiating oligodendrocytes (Sun et al., 2018; Zhang et al., 2014). However, the upstream signaling pathways leading to upregulation and downregulation, as well as activation of this pathway remain unclear. Axonal biophysical properties (Goebbels et al., 2017; Lee et al., 2012; Mayoral et al., 2018; Voyvodic, 1989; Bonetto et al., 2020), neuronal activity (Gyllensten and Malmfors, 1963; Tauber et al., 1980; Demerens et al., 1996; Lundgaard et al., 2013; Hill et al., 2014; Gibson et al., 2014; Gautier et al., 2015; Mensch et al., 2015; Mitew et al., 2018; Ortiz et al., 2019; Bacmeister et al., 2022; Bonetto et al., 2020), and growth factor release (Geraghty et al., 2019; Lundgaard et al., 2013) have all been shown to modulate myelination. Whether and how these cues directly interact with pro apoptotic machinery to regulate oligodendrocyte integration and myelination remains an open question.

If procaspase-3 is upregulated to allow for a rapid fate decision between death and integration, and if this is important to control the temporal sequence of myelination, why do neonatal differentiating oligodendrocytes lack high procaspase-3 levels? There are several conceivable explanations, including: neonatal BCAS1+ procaspase-3-cells are 1) OPCs, 2) an intermediate stage between OPCs and differentiating oligodendrocytes, 3) all dying through cleaved caspase-3-mediated apoptosis, 4) cannot die through cleaved caspase-3-mediated apoptosis, or 5) stalled as differentiating oligodendrocytes. Our findings suggest that we can eliminate the first two possibilities, as neonatal BCAS1+ procaspase-3-cells are likely *bona fide* differentiating oligodendrocytes since they lack OPC markers and are labeled by several differentiating oligodendrocyte markers.

Furthermore, our data do not support the third possibility, as we rarely detected cleaved caspase-3+ BCAS1+ cells in the neonatal cortex and corpus callosum, in line with previous work showing very few CC1+ cleaved caspase-3+ cortical cells at P10 (Hill et al., 2014) or CNP-EGFP+ cleaved caspase-3+ cells at P12 in the cortex and corpus callosum (Guardia Clausi et al., 2012). These low numbers are difficult to reconcile with previous observations suggesting that 20% of premyelinating oligodendrocytes appear to be fragmented and dying in the P7 and P11 cortex (Trapp et al., 1997), but this discrepancy might result from different detection methods for dying cells (cleaved caspase-3 labeling vs. a fragmented morphology). Nonetheless, our data and these previous observations are in agreement that the majority of neonatal differentiating oligodendrocytes are not dying.

This suggests a fourth possibility: differentiating oligodendrocytes rarely die through cleaved caspase-3-mediated apoptosis in the neonatal forebrain. Myelination has been suggested to occur in two modes, an activity-dependent mode and an activity-independent mode (Lundgaard et al., 2013; Koudelka et al., 2016), and these have been proposed to correlate with developmental and adaptive or experience-dependent myelination (Monje and Káradóttir, 2021; Bechler et al., 2018). It is conceivable that during early development, differentiating oligodendrocytes do not upregulate high levels of procaspase-3, as baseline myelination proceeds unimpaired and thus, there is no fate decision to be made between death or integration. However, once developmental myelination is complete, myelination may become activity-dependent or adaptive, and differentiating oligodendrocytes may be faced with a fate decision: to integrate and myelinate, or to die. Procaspase-3 upregulation may therefore be a mechanism to allow for this fate decision to occur should the right environmental signals promoting integration not be received.

On the other hand, our data that blocking caspase-3 reduced the number of CNP+ oligodendrocytes rather suggest that increased procaspase-3 promotes oligodendrocyte differentiation. Although it is mainly studied as an apoptotic executioner, caspase-3 has also been shown to promote cell survival, proliferation, and differentiation in various tissues and cell types (Eskandari and Eaves, 2022). For instance, blocking or knocking out caspase-3 in neurospheres delays neural stem cell differentiation into neurons, oligodendrocytes, and astrocytes without altering apoptosis (Fernando et al., 2005). Of note, non-apoptotic functions have been attributed to both cleaved caspase-3 and procaspase-3 (Eskandari and Eaves, 2022), with a recent study suggesting that the pro domain alone can promote survival and proliferation of human cells in vitro (Eskandari et al., 2024). This could potentially explain why we found that a reduction in procaspase-3+ cells with limited change in cleaved caspase-3+ cells resulted in fewer CNP+ positive oligodendrocytes.

A role for procaspase-3 in differentiation would support the fifth possibility, that neonatal BCAS1+ CC1+ procaspase-3-cells could be stalled in the differentiating oligodendrocyte stage. The idea of stalled differentiating oligodendrocytes was recently proposed in the context of de novo myelination following learning, where differentiating oligodendrocytes are detected within hours of motor learning, while new myelin is detected weeks after learning (Bonetto et al., 2021), even though a study in zebrafish suggests that myelination by individual oligodendrocytes happens within hours, not days (Czopka et al., 2013). Stalled differentiating oligodendrocytes in neonates could be consistent with the presence of BCAS1+ CC1+ cells as early as P4, but the first MBP+ myelin sheaths only being detected at P10 (Vincze et al., 2008). Moreover, the time of onset of procaspase-3 upregulation in the corpus callosum correlates with the onset of rapid myelination (Hamano et al., 1996, 1998). If procaspase-3 upregulation does play a role in differentiation, it will be interesting to determine in future studies whether it is the zymogen itself or cleaved caspase-3 that promotes differentiation, how it does so, and which intrinsic or extrinsic signals mediate procaspase-3 upregulation in differentiating oligodendrocytes and subsequent downregulation in mature cells. A further outstanding question is whether the upregulation of procaspase-3 and other pro apoptotic factors such as PUMA (Sun et al., 2018) renders differentiating oligodendrocytes more sensitive to cytotoxic insults. Answering these questions will provide additional insights into differentiating oligodendrocyte resilience and potential targets to promote myelination in degenerative conditions.

## ACKNOWLEDGMENTS

We thank members of the Hill lab for helpful discussions and feedback on the project. This work was supported by the National Institutes of Health R01NS122800 and the Esther A. & Joseph Klingenstein Fund and Simons Foundation to R.A.H. and a Postdoctoral Fellowship FG-2307-42173 from the National Multiple Sclerosis Society to Y.K.

## AUTHOR CONTRIBUTIONS

Y.K. and R.A.H. conceived and designed the study. Y.K. performed all the experiments and most of the data analysis and quantification. T.W.C. prepared the P60 tissue and contributed to cell quantification in Figure 1. E.T.P. contributed to cell quantification in Figure 1. R.A.H. and M.E.C. prepared slice cultures. Y.K. and R.A.H. wrote the paper and R.A.H. supervised the study.

## METHODS

### Animals

All animal procedures were submitted to and approved by the Institutional Animal Care and Use Committee at Dartmouth College. The initial characterization of procaspase-3 was performed in 8-week-old C57BL/6 mice purchased from the Jackson Laboratory. To examine procaspase-3 density with age (Figures 4 and 5), stock mice from various transgenic lines were used between P4 and P737, as specified in the text. All mice were bred on a C57BL/6 background. Mice were housed in a 12 hours light/dark cycle in an animal vivarium with controlled temperature (22 °C) and humidity (30–70% relative humidity). Food and water were provided ad libitum.

### Immunohistochemistry

P14 to P737 mice were deeply anaesthetized with ketamine (100mg/kg) and xylazine (10mg/kg) and perfused with ice-cold PBS followed by 25-30 mL of 4% paraformaldehyde (PFA). Brains were dissected from the skull and postfixed in 4% PFA overnight at 4 °C, except for NG2 staining when brains were postfixed in 4% PFA for one hour at room temperature (RT). Brains from P4 to P10 mice were dissected immediately after euthanasia and fixed in 4% PFA for 24 hours. 75 to 100 μm-thick sections were cut on a vibrating microtome (Leica VT 1000S) and stored at -20 °C in cryostorage solution composed of 1% Polyvinylpyrrolidone, 30% sucrose, and 30% ethylene glycol in 0.2 M sodium phosphate buffer (pH 7.4).

For immunolabeling, sections were washed three times in PBS for 10 minutes. For stainings shown in Figures 4 and 5, sections were then incubated in a 10 mM copper(II) sulfate, 50 mM ammonium acetate solution (pH 5) for one hour at RT on a rotating shaker to quench endogenous fluorophores and lipofuscin (Schnell et al., 1999). Next, sections underwent antigen retrieval, where they were incubated for 2 minutes at 98°C in preheated 10 mM Tris, 1 mM EDTA, and 0.05% Tween 20 in PBS (pH 9). Once sections had cooled down to RT, they were incubated with primary antibodies in blocking and permeabilization solution (1% BSA, 0.3% Triton X 100 in PBS) overnight at RT on a rotating shaker. The following primary antibodies were used: goat anti-Human/Mouse Caspase-3 (R&D Systems, Cat# AF-605-SP, 1:100-1:200), guinea pig anti-CNP1 (Synaptic Systems, Cat# 355 004, 1:500), mouse anti-CNPase (Biolegend, Cat# 836404, 1:750), rabbit anti-OLIG2 (EMD Millipore, Cat# AB9610, 1:500), rabbit anti-PDGFRA (Cell Signaling, Cat# 3174S, 1:500), goat anti-PDGFRA (R&D Systems, Cat# AF1062, 1:1000), guinea pig anti-BCAS1 (Synaptic Systems, Cat# 445 004, 1:750), mouse anti-CA II (Santa Cruz, Cat# 48351, 1:500), rat anti-CA II (R&D Systems, Cat# MAB2184, 1:200), mouse anti-CC1 (Millipore, Cat# MABC200, 1:300), rabbit anti-cleaved caspase-3 (Cell Signaling, Cat# 9661S, 1:700), rabbit anti-NG2 (Millipore, Cat# AB5320, 1:500), and rabbit anti-TCF7L2 (Cell Signaling, Cat# 2569, 1:500). The following day, sections were washed three times in PBS for 10 minutes. Sections were incubated with secondary antibodies in blocking and permeabilization solution for one hour at RT on a rotating shaker. Secondary antibodies were conjugated to Alexa Fluor 488, 555, or 647 (ThermoFisher or Jackson ImmunoResearch, 1:500), or CF660C (Biotium, 1:500). Sections were then washed twice in PBS for 10 minutes and incubated with 1:2000 Hoechst 33342 (ThermoFisher, Cat# H3570) in PBS for 20 minutes at RT on a rotating shaker. Following a final 10 minutes wash in PBS, sections were mounted on glass slides with ProLong Diamond Antifade Mountant (ThermoFisher, Cat# P36970).

1:100 goat anti-Human/Mouse Caspase-3 was used for images presented in Figures 1, 2A (with OLIG2, PDGFRA and BCAS1 labeling),B (with CNP labeling),E, 3A,B,E, and 4; and the corresponding quantifications in Figures 1C,D, 2C,D, and 3D. 1:200 goat anti-Human/Mouse Caspase-3 was used in all other instances.

### Slice culture

Hippocampal slices were prepared from P9 mice, as previously described (Hill et al., 2013; Sherafat et al., 2018). Briefly, 300μm-thick coronal forebrain slices were cut with a vibrating microtome (Leica VT1000S or Precisionary Instruments VF-310-0Z) and hippocampi were micro dissected in ice-cold dissection medium comprised of 124 mM NaCl; 3 mM KCl; 1.24 mM KH_2_PO_4_; 4 mM MgSO_4_; 2 mM CaCl_2_; 26 mM NaHCO_3_; 10 mM D-glucose; 2 mM ascorbic acid; and 0.075 mM adenosine. The slices were placed on Millicell culture inserts with 0.4 μm pore size (Millipore, Cat# PICMORG50) and kept in a humidified 37 °C, 5% CO_2_ incubator. The culture medium was comprised of 50% Minimal Essential Medium with Earle’s Salts; 25% HBSS without calcium chloride, magnesium chloride, or magnesium sulfate; 25% horse serum; 25 mM HEPES; 0.4 mM ascorbic acid; 1 mM L-glutamine; and 1 mg/L insulin. pH was adjusted to 7.2 with 1 M NaOH. The culture medium was changed the day after dissection, and every two days afterwards. On day in vitro (DIV) 7, the slices were treated with either 20μM Z-DEVD-FMK (Chen et al., 2013; Kanthasamy et al., 2006; Nishida et al., 2012) or the vehicle only.

For immunostaining, slices were fixed in 4% PFA for 30 minutes on DIV 14 and washed in PBS. Membranes were cut out of the inserts and staining was performed as described above.

### Imaging and quantification

Images were acquired on an upright laser scanning microscope (Leica SP8) equipped with a 10X air objective (Leica NA 0.4), a 20X air objective (Leica NA 0.75) and a 63X oil objective (Leica NA 1.4). Fluorophores were excited with the following laser lines: 405 nm, 488 nm, 552 nm and 633 nm. Multichannel images were acquired sequentially, from longest to shortest wavelength.

For Figures 1, 2, 4, 5, three non-overlapping images of the somatosensory cortex or two images of the corpus callosum were taken from two brain slices per mouse and the number of cells was counted in Fiji (Schindelin et al., 2012). For Figure 3 and Figure 5f, six non-overlapping images of the somatosensory cortex from two slices per mouse were used to increase sampling of rare cell populations. Somatosensory cortex images were acquired between the pial surface and ∼500 μm below. Corpus callosum images were acquired at the midline, between Bregma and Bregma -1.00 mm. Cells were counted within a fixed volume, which was used to calculate cell densities in Figures 4-5. The experimenter was blinded to age for Figures 4-5. For Figure 6, six non-overlapping images per slice were used to count cells within a fixed volume and calculate densities. The experimenter was blinded to the treatment group.

### Statistical analysis

Data are shown as mean ± standard error of the mean (s.e.m.) and *n* numbers are indicated in the figure legends. Statistical analyses were performed with GraphPad Prism. When comparing two groups, unpaired two-tailed t-tests were computed, and variance was tested with an F-test.

When variance was unequal, Welch’s corrected t-test was used. When comparing three or more groups, a one-way ANOVA was used; variance was tested with a Brown-Forsythe test and Welch’s ANOVA was used when variance was unequal. Šidák’s (equal variance) or Dunnett’s T3 (unequal variance) pairwise multiple comparisons were used.

